## Supplementary Materials for "Goal-directed vocal planning in a songbird"

### Supplementary Materials for:
### Goal-directed vocal planning in a songbird

Anja T. Zai, Anna E. Stepien, Nicolas Giret, Richard H.R. Hahnloser

**Recovery of baseline after pitch reinforcement but not after duration reinforcement**

Zebra finches learn both syllable pitch and syllable duration from their tutors (1,2) but are these distinct sound features equally well maintained in adulthood? Both pitch and duration can be driven away from baseline using aversive reinforcers (3). To test whether recovery of syllable duration is similarly fast as recovery of pitch (4), we induced birds to shorten or lengthen a targeted song syllable by playing a loud WN stimulus when the syllable duration was shorter or longer than a manually set threshold. However, we found that unlike for pitch, after withdrawal of reinforcement, these WNdur birds did not reliably recover their baseline syllable duration: after 4 days of song practice without reinforcement, their average normalized residual syllable duration ($NRD$) — the remaining fraction of duration difference to baseline since release — was significantly different from zero ($NRD = 42\%, p = 0.007$, two-sided t-test for $NRD = 0$, $N = 12$ birds, Suppl. Fig. 1), suggesting that adult birds do not maintain syllable duration as rigidly as they do maintain pitch.

Indeed, in contrast, pitch-reinforced (WNC) birds fully recovered baseline pitch 4 days after withdrawal of reinforcement (average $NRP = 1\%, p = 0.91$ two-sided t-test for $NRP = 0$, $N = 18$ birds, Supplementary Fig. 1 B, C, same data as in Fig. 1 H, I). Moreover, the recovery of pitch was more extensive than that of duration: after 4 days the average $NRP$ was smaller than the average $NRD$ ($p = 0.001$, tstat $= -3.56$, $df = 28$, two-tailed t-test). The difference between $NRP$ and $NRD$ persisted over time and was still present after 10 days of practice ($NRP = -2\%$ and $NRD = 33\%$ after 10 days of practice, $p = 0.04$, $t$stat $= -2.26$, $df = 18$, $N = 10$ WNC and $N = 10$ WNdur birds recorded 11 days after WN, two-tailed t-test), showing that syllable duration is not recovered as reliably as pitch after withdrawal of white-noise reinforcement.

WNC birds received WN reinforcement during $11 \pm 3$ days on average (range $8 - 15$ days) and during this period they changed their pitch by $0.24 \pm 0.22$ $d'$/day (total shift $2.5 \pm 1.8$ $d'$) on average, which did not significantly differ from the behavior of WNdur birds ($p = 0.44$, tstat $=$ $0.78$, $df = 28$, two-tailed t-test) that received WN reinforcement during $10 \pm 5$ days on average (range $5 - 23$ days) and changed syllable duration during this period by $0.36 \pm 0.17$ $d'$/day (total shift $2.9 \pm 0.7$ $d'$) on average ($p = 0.14$, tstat $= -1.49$, $df = 28$, two-tailed t-test). Furthermore, we did not find a correlation between the number of days with reinforcement and the normalized residual pitch/duration after 4 days of practice in either WNC birds ($R = 0.20, p = 0.43$), WNdur birds ($R = 0.05, p = 0.88$), or in both groups combined ($R = -0.01, p = 0.94$), nor between the total shift and the normalized residual pitch/duration after 4 days of practice in either WNC birds ($R = 0.34, p = 0.17$), WNdur birds ($R = -0.12, p = 0.71$), or in both groups combined

($R = 0.18$, $p = 0.34$), suggesting that the difference in recovery behavior between WNdur and WNC birds is neither due to differences in time spent with reinforcement nor to differences in learning speed, but due to differences in attachment to these sounds features.

This difference in recovery behavior is consistent with the literature. Ali et al. (3) found that pitch recovers at roughly half the rate it changed during reinforcement (12.1 Hz/day during recovery vs. 26.0 Hz/day during reinforcement). By contrast, they found that duration (of mostly syllables+gaps) reverted towards baseline at only about a quarter of the rate measured during reinforcement (0.8 ms/day vs. 3.4 ms/day) (3), suggesting that recovery of duration proceeds at a much slower pace than birds are capable of. Similarly, Roberts et al. found pitch recovered faster than it changed during reinforcement whereas syllable+gap durations recovered slower than they changed during reinforcement (5 birds each) (5). Both studies used only few birds and did not directly compare the recovery of duration versus pitch. Also, the published reports preclude a comparison in terms of the unit-free $d'$ measure post-hoc because the baseline pitch and the duration variability are not reported in these studies.

Interestingly, in the context of Figure 4, duration reverts similarly to pitch. After 4 days of practice WNdur birds showed a significant reversion towards baseline ($d' = -2.03$, $p = 5.5 * 10^{-4}$, tstat $= -4.37$, $N = 11$ WNdur birds, one-sided t-test) indicating that WNdur birds are capable of revertive changes beyond the planning limit ($d' \simeq 1$) found for pitch reversion. Thus, it remains unclear whether the difference in NRP and NRD recovery (Fig. S1) could be confounded by the disparity in experimental parameters (i.e. difference in total shifts). Nevertheless, all studies (3,5) and our analysis concur that the recovery of baseline pitch is more efficient than the recovery of duration, suggesting that duration is less accurately maintained than pitch. We therefore decided to perform all experiments using pitch as the targeted song feature and not syllable duration.

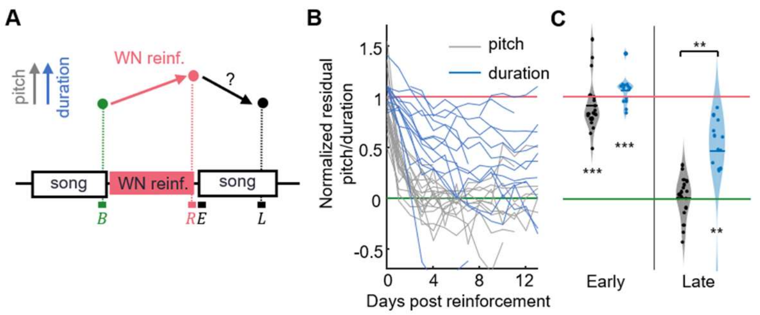


**Fig. S1. Birds rapidly recover pitch but not duration after reinforcement learning.** (**A**) Birds were pitch or duration reinforced using WN. Pitch recovery from the reinforced ($R$) state towards the baseline ($B$) target is evaluated in an early (E, no practice) analysis window and a late ($L$) window 4 days later. (***B***) After ending WN reinforcement on day 0, pitch (gray lines, *N=19* experiments from 18 birds) recovered their NRP/NRD faster and more fully than duration (blue lines, N=14 experiments from 13 birds). (**C**) Violin plots of same data restricted to early and late analysis windows (****p<0.001*, ***p<0.01*, two-tailed t-test of $NRP = 0$).

| **Group** | **No practice (early)** | | | | **5 days practice (late)** | | | |
|---|---|---|---|---|---|---|---|---|
| **Reference time** | $R$ =last 2 h of reinforcement | | $R$ =2 h time-of-day aligned | | $R$ =last 2 h of reinforcement | | $R$ =2 h time-of-day aligned | |
| | d’ | p | d’ | p | d’ | p | d’ | p |
| **WN deaf** | -0.6 ±0.3 | 0.007 | -0.47 ±0.3 | 0.03 | -1.3 ±0.2 | $p<10^{-4}$ | -1.1 ±0.2 | $p<10^{-4}$ |
| **Deaf LO** | -0.09 ±0.13 | 0.25 | 0.34 ±0.07 | p>0.99 | -0.17 ±0.13 | 0.10 | 0.35 ±0.11 | p>0.99 |

**Table S1: Bootstrapped pitch differences between reinforced and deaf control birds.** The reported p values correspond to the null hypothesis that release from reinforcement causes no pitch change (bootstrapped, see methods). Two separate analysis windows $R$ are considered: 1) the last 2 h of reinforced song on the evening before, and 2) a time-of-day matched 2-h window on the morning before (discounting for circadian trends). WNd birds changed pitch towards baseline, but dLO birds did not.
